# Label-Free Fluorescence Microscopy Reveals Multiphase Organization in Biomolecular Condensates

**DOI:** 10.1101/2025.07.13.663051

**Authors:** Bibek Acharya, Sean C. Castillo, Rukhillo Kodirov, Anisha Shakya

## Abstract

Phase transitions of proteins and nucleic acids (NA) leading to the formation of biomolecular condensates have been linked to various biological functions. Given the growing number of proteins/NA predicted to undergo liquid-liquid phase separation (LLPS), efficient tools to investigate this behavior are critical to advancing our understanding of biomolecular condensate function. The current standard used to study LLPS involves techniques that utilize exogenous fluorophore labels. The labeling process is often costly and time-consuming and comes with associated complexity that arises from unknown interactions from the bulky fluorescent tags. These aspects limit high throughput analysis of protein/NA phase separation based on external fluorophore labeling. Here, we report the discovery that intrinsic fluorescence, well into the visible spectrum, arises as an emergent property of biomolecular condensates. Leveraging this intrinsic fluorescence, we study condensate formation, directly measure their internal dynamics via Fluorescence Recovery after Photobleaching (FRAP), and examine the 3D morphology and transitions to various multiphase architectures. Through this approach, we find that a variety of G-quadruplex DNA readily form droplets with histone H1 and display dynamic exchange. In addition, we directly demonstrate that the 3D morphology, core-shell architecture, and sub-compartmentalization of condensate droplets are tunable via the charge ratio of components in solution and NA hybridization. Our method utilizes an inherent property of condensates, thus is broadly applicable to any phase-separated systems and can advance our understanding of biological phase transition.

## Introduction

Within the cell are several structures and organelles whose physical boundary is not defined by the presence of a lipid membrane. Liquid-liquid phase separation (LLPS) has emerged as an important mechanism to explain the existence of membrane-less cellular compartments. LLPS allows for the concentration of certain protein and nucleic acids (NA) into liquid-like compartments termed biomolecular condensates (1, 2). The emergent properties of biomolecular condensates have been linked to various regulatory cellular processes (2-7) and as such, the number of studies on the condensate properties of proteins and NAs have significantly increased in recent years. Accessing characteristics such as morphology, merging dynamics, internal diffusion, and dependence on external perturbations typically involves fluorescence imaging of biomolecular condensates (8, 9).

Condensate components are often labeled with bulky fluorescent tags containing aromatic chromophores due to the widespread use of fluorescence techniques as the primary method of investigation. A major drawback of exogenous fluorescence labeling, beyond being expensive and time-consuming, is the unknown effect of the fluorescent tags on the native structure and local interactions of the labeled protein or NA. Development of label-free techniques which yield efficient characterization of biomolecular condensates is therefore highly desirable. In this study, we identify an emergent property of biomolecular condensate that allows fast, label-free imaging and spectroscopy of the condensates well into the visible spectrum. We leverage this property to directly study phase behavior, dynamics, and modulation of 3D morphology and multiphase architecture of biomolecular condensates.

Typically, photonic emissions of luminescent molecules are attributed to fluorescence from local excitation of highly conjugated molecules. However, research in the early 2000s demonstrated non-conjugated molecules, optically inactive in isolation, could exhibit strong absorption and fluorescence in the UV and visible regions of the spectrum upon aggregation (10-12). While the mechanism of this phenomenon is still unclear, several possibilities have been proposed, including aggregation/clustering induced emission (AIE) (12). For aggregates of non-conjugated molecules, fluorescence is most often reported in the blue region of visible spectrum upon near-UV excitations. For nucleic acids, absorption and fluorescence in the blue region of the visible spectrum is detectable (13, 14), though significantly weaker than absorption in the UV range at roughly 250 nm. As condensation results in a highly localized concentration of interacting biomolecules (1, 2), we hypothesize condensate formation causes intrinsic characteristic absorbance and fluorescence in the visible spectrum from the condensates of nucleic acids. For both protein-DNA and polypeptide condensates we report strong intrinsic absorption and fluorescence in the middle of the visible spectrum (λ_ex_ ∼ 532 nm, λ_fl_ ∼ 550 nm), though the emission intensity increases significantly in presence of DNA.

Many factors contribute to biological phase transitions. To fully understand the determinants, detailed in vitro studies are important. As inferred from such studies, critical concentrations and fine-tuning of components are required for LLPS. The phase behavior of a condensate can be tuned by varying parameters such as salt concentration, pH, temperature, as well as molecular structural details (1, 15, 16). Until now, label-free systems commonly required bright-field imaging or turbidity measurements to detect and characterize phase transitions. These methods often do not fully show the accurate quantitative picture as they cannot capture condensate morphology. Many biomolecular condensates observed in cells have intricate architectures, such as coexisting sub-compartments or core-shell organization (2, 17, 18). Factors determining such spatiotemporal organization is not yet understood. These varied architectures can result from interactions that arise due to the varied secondary and tertiary structures of the proteins and nucleic acids involved. Here, we examine the innate fluorescence as an emergent property across a wide assembly of nucleic acid condensates.

We study the condensates of histone and G-quadruplex DNA, condensates of oppositely charged polypeptides, as well as DNA-polypeptide condensates through detection of their intrinsic fluorescence. Coupling our imaging scheme with fluorescence recovery after photobleaching (FRAP), we find that the internal dynamics of histone-G-quadruplex is strongly dependent on the structure of the G-quadruplex. The observation that internal dynamics are strongly dependent on the molecular structure of the phase separating components is further observed in poly(R-lysine)-poly(L-glutamic acid) and poly(R-lysine)-poly(L-aspartic acid) condensates, which display a difference in internal diffusion by more than an order of magnitude.

Our approach allows us to discern the 3D spatial organization of components in multiphase condensates, without the need of extensive labeling by fluorescent tags. We find that by tuning the ratio of positively and negatively charged species through the adjustment of the ratio of net positive charge from protein amino acids to DNA phosphate (N:P), the 3D morphology of condensates can be modulated. Additionally, we show that NA hybridization drives core-shell architecture of condensates in an N:P ratio dependent manner. Finally, we demonstrate that the differences in fluorescence intensities arising from protein-rich vs NA-rich condensates can be employed to detect the different phases in multiphase condensate.

## Results

### Fluorescence emission from DNA-protein condensates in absence of fluorophore labeling

To test whether intrinsic fluorescence emission is observed in the visible spectrum upon LLPS, we mixed solutions of the homopolypeptide, poly(L-lysine) (n=240, PLL_240_), and a 22 nucleotide (nt) single-stranded (ss) DNA, poly(A)_22_, which have been previously shown to form phase-separated droplets together under a variety of conditions (18). Upon excitation at 532 nm, clear fluorescence emission from the LLPS droplets is observed at 550-750 nm, as evidenced from the confocal fluorescence microscopy images of PLL_240_-poly(A)_22_ droplets (**Figure 1a**).

**Figure 1.**
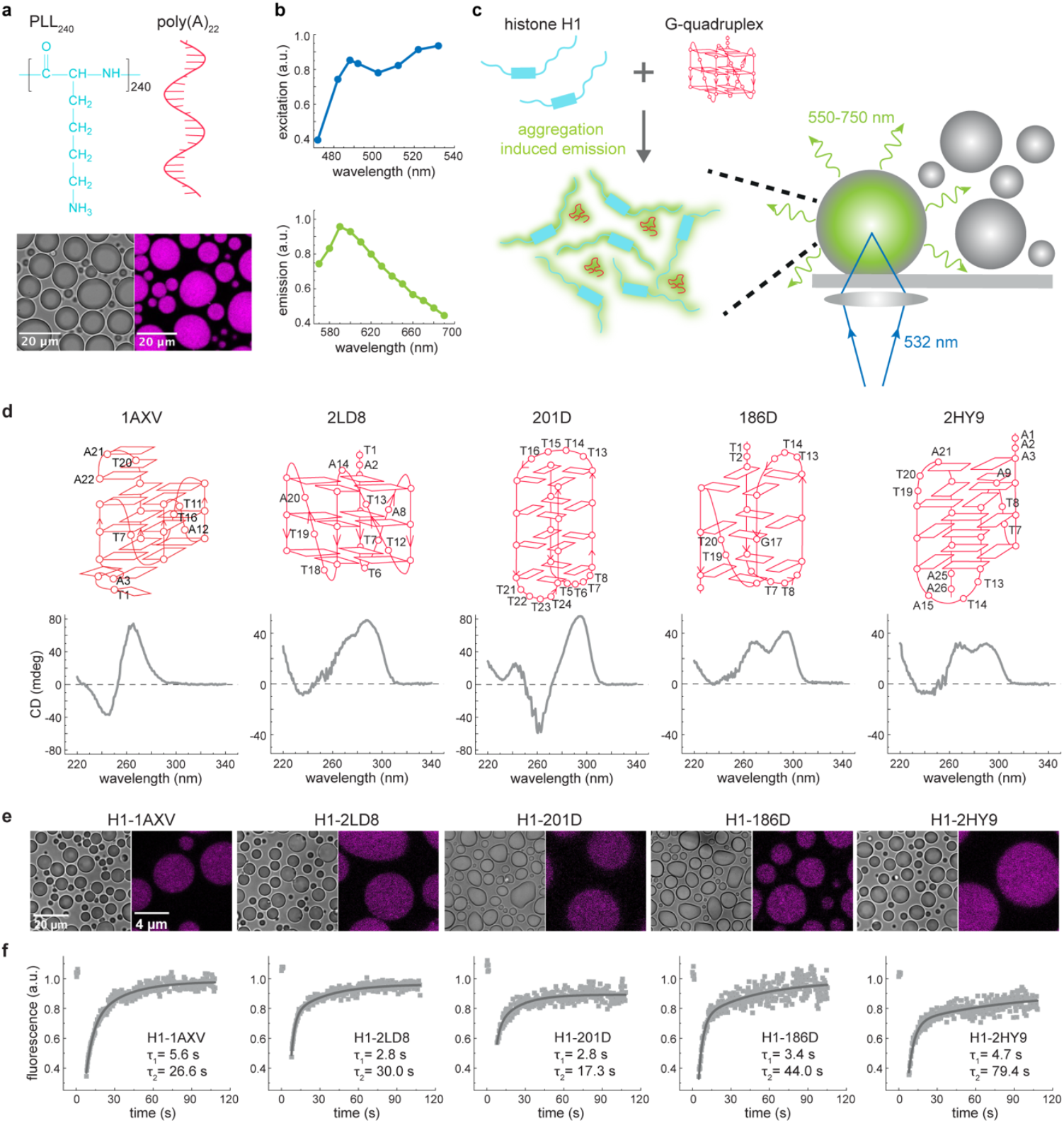
Detection of visible wavelength intrinsic fluorescence reveals DNA condensate formation and internal dynamics. (**a**) Intrinsic fluorescence detected upon liquid-liquid phase separation of poly-L-Lysine (PLL_240_) and single stranded DNA (poly(A)_22_). (**b**) Characterization of intrinsic fluorescence of PLL_240_-poly(A)_22_ droplets by measuring excitation (top) and emission (bottom) spectra reveals that the fluorescence can be detected well-into the visible wavelength range upon condensate formation. (**c**) Schematic depicting fluorescence emission upon condensing the histone protein, H1 and G-quadruplex DNA into LLPS droplets, potentially via aggregation induced emission. Confocal images were obtained using 532 nm excitation in absence of fluorophore tags. Emission was detected using 550-750 nm bandpass filter. (**d**) Folding topology of five different G-quadruplexes (38) used in this study (top) and their corresponding circular dichroism (CD) spectra (bottom). (**e**) Bright-field (left) and confocal (right) images of droplets of five different G-quadruplexes with H1 (buffer: 10 mM phosphate, pH 7.4, 0.2 mM EDTA, 50 mM KCl). (**f**) Fluorescence recovery after photobleaching (FRAP) experiments on the droplets shows photo-bleaching and near complete recovery (see **Figure S1**). The FRAP curves can be fit to two exponentials with the decay times varying with the G-quadruplex.

To characterize the fluorescence arising upon LLPS, we obtained the excitation and emission spectrum of the PLL_240_-poly(A)_22_ droplets. The excitation spectrum shows broad absorption with maxima at 488 nm and 532 nm (**Figure 1b, top**). The fluorescence emission spectrum (532 nm excitation) shows a large Stokes shift of ∼50 nm, with emission maximum at 590 nm (**Figure 1b, bottom**). Native protein fluorescence is typically attributed to the presence of aromatic amino acids (tryptophan, tyrosine, and phenylalanine) with fluorescence absorption and emission reported at wavelengths <400 nm (19). Clear excitation and emission well into the visible spectrum upon LLPS of non-aromatic polypeptides, lead us to suspect this intrinsic fluorescence emission is a general property of any biomolecular condensate system.

We next utilized condensate intrinsic fluorescence to study the phase transitions of the chromatin architectural protein, histone H1, in presence of a variety of G-quadruplex DNA (**Figure 1c, d**). LLPS of histone H1 is shown to be important for organization of genomic DNA into gene-silenced regions (heterochromatin) (16, 20, 21). Previously we have shown that H1 undergoes LLPS to form liquid-like droplets in presence of ssDNA, double stranded (ds) DNA, and nucleosome arrays of various lengths (15, 16). Here we report H1 also forms LLPS droplets with various G-quadruplex DNA (**Figure 1d-e**). The droplets readily form under conditions of 10 mM phosphate buffer with 50 mM KCl, 11 µM H1, and 20-25 µM DNA concentration (resulting N:P ratio ∼1). As expected, upon excitation at 532 nm, clear fluorescence emission was detected between 550-750 nm from the H1-G-quadruplex droplets (**Figure 1e**).

By examining droplet intrinsic fluorescence, we were able characterize the internal dynamics of the H1-G-quadruplex droplets using FRAP (**Figure 1f, S1**). Though this fluorescence cannot be uniquely assigned to a specific component, we find the local intensity can be effectively bleached and dynamically recovered from the exchange of biomolecules. After applying this approach to droplets formed from six different G-quadruplex DNA of 20-28 nucleotide lengths, the observed FRAP exchange rates varied based on the respective G-quadruplex (**Figure 1f**). As fluorescence was detected without the use of exogenous fluorescent tags, the recovery of fluorescence directly reflects the dynamics of the biomolecules within the condensates.

### Intrinsic fluorescence emission from non-DNA condensates

In the H1-G-quadruplex droplets, the excitation and fluorescence (532 nm and 550-750 nm, respectively) was observed at much longer wavelengths compared to previously reported aggregation induced emission (AIE) from protein aggregates (22, 23). To test if intrinsic fluorescence only arises in LLPS systems containing DNA, we next studied the LLPS of polypeptides of non-conjugated amino acids in absence of DNA.

After mixing oppositely charged homopolypeptides in solution at N:P ∼1, droplets formed via LLPS driven by interactions of the positively and negatively charged polypeptide chains. **Figure 2** shows droplets formed after mixing the positively charged polypeptide poly(R-lysine), PRK_100_, with the negatively charged polypeptides poly(L-glutamic acid), PLE_100_, or poly(L-aspartic acid), PLD_100_. We find that droplets of these polypeptides in absence of DNA also fluoresce upon 532 nm excitation, though the emission is significantly weaker compared to emission from droplets containing DNA.

**Figure 2.**
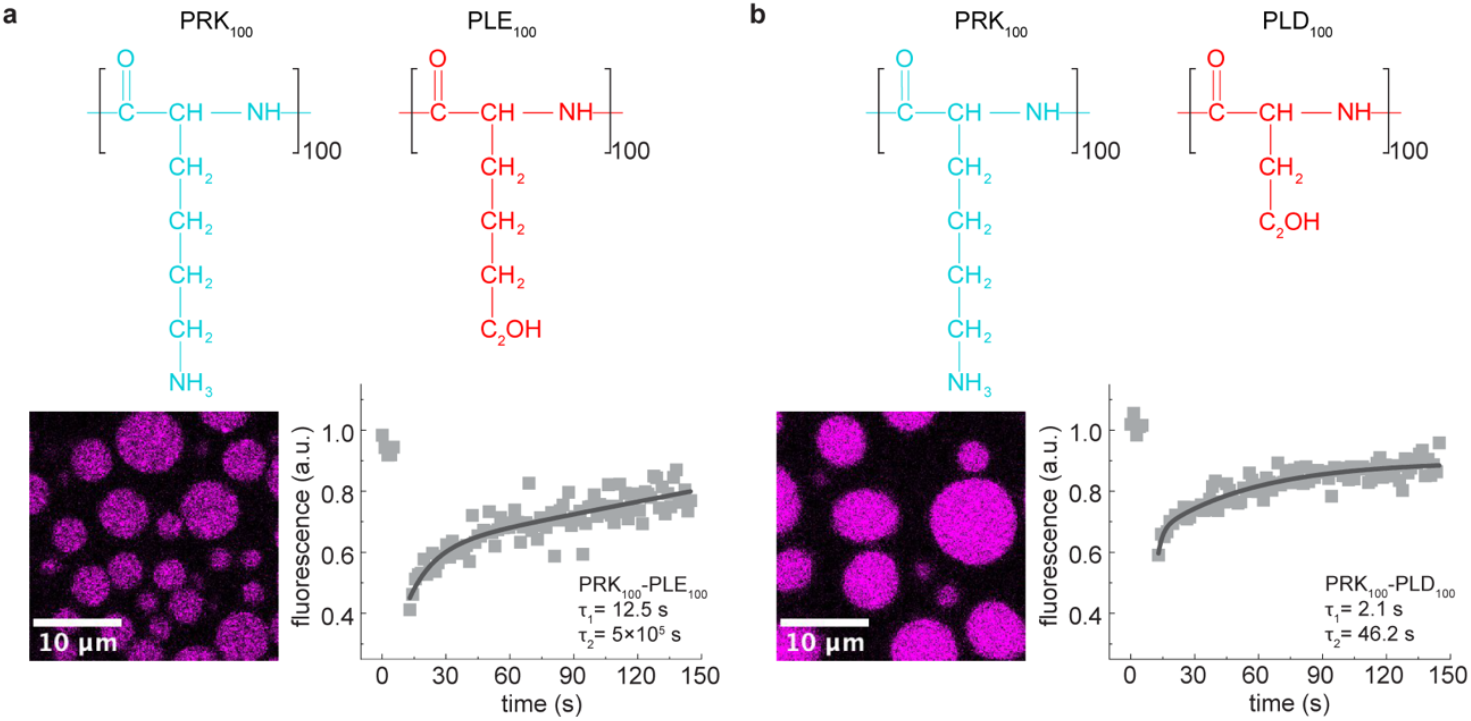
Visible wavelength intrinsic fluorescence can be detected in absence of DNA. (**a**) Intrinsic fluorescence detected upon LLPS of the oppositely charged polypeptides, poly-R-Lysine (PRK_100_) and poly-L-glutamic acid (PLE_100_) (bottom left). FRAP experiments on the PRK_100_-PLE_100_ droplets shows photo-bleaching and near complete recovery (bottom right). (**b**) Intrinsic fluorescence detected upon LLPS of the oppositely charged polypeptides, poly-R-Lysine (PRK_100_) and poly-L-aspartic acid (PLD_100_). FRAP experiments on the PRK_100_-PLD_100_ droplets shows photo-bleaching and near complete recovery. Excitation and fluorescence emission were at the same wavelengths as that for DNA-containing condensates depicted in **Figure 1**.

The internal dynamics of the polypeptide droplets were assessed using intrinsic FRAP (**Figure 2a-b, bottom panels**). The fast recovery rates after photobleaching indicate rapid diffusion of the macromolecules within the condensates, as expected in a dynamic liquid-like condensate. In addition, PRK_100_-PLE_100_ and PRK_100_-PLD_100_ droplets show significant differences in the FRAP recovery rates indicating the intrinsic fluorescence are governed by properties of the individual droplet system, consistent with the observation for protein-DNA droplets (**Figure 1**).

### Droplet morphology can be tuned by DNA concentration-dependent overall charge ratio

Bright-field imaging or turbidity measurements are often the common way to detect and characterize phase transitions in absence of fluorophore tags. However, these methods do not give the quantitative picture as they cannot fully account for the morphology of resulting condensates. Detection of condensate intrinsic fluorescence overcomes this limitation, as 3D video-microscopy of condensates allows for more accurate characterization of phase behavior. Using this approach, we can directly demonstrate the dependence of condensate morphology on the overall charge composition of components in solution.

For mixtures of charged polypeptides/proteins and nucleic acids, it is useful to represent the concentration ratio in terms of the overall charge ratio, designated as N:P, between the components. **Figure 3a** shows LLPS droplets formed after mixing PLL_240_ and poly(A)_22_ at solution N:P ratios ranging from 0.2 to 3.5. As seen in 3D confocal images, circular droplets are formed for all tested N:P ratios with droplet fluorescence intensity decreasing with increasing N:P. Interestingly, the N:P ratio modulates the contact angle of the droplet and the surface (**Figure 3a bottom panel**, x-z plane), with the contact angle decreasing upon increasing N:P.

**Figure 3.**
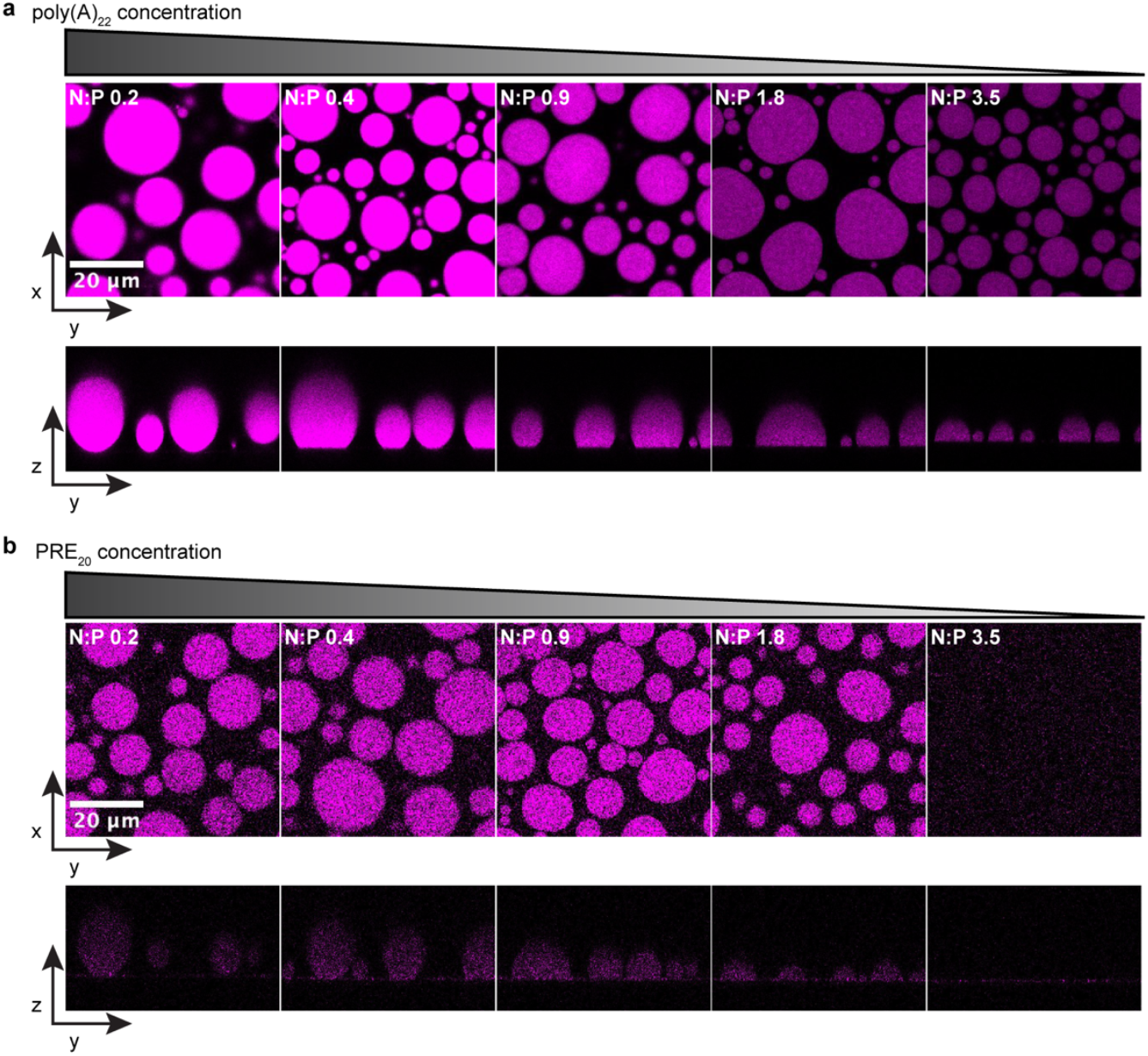
Droplet 3D morphology is tuned by DNA concentration-dependent overall charge ratio. (**a**) Confocal intrinsic fluorescence images of PLL_240_-poly(A)_22_ droplets at increasing N:P (ratio of net positive charge to negative charge of macromolecules in solution) in the x-y plane (top panels) z-y plane (bottom panels). Droplets are more elevated in the z-axis at lower N:P at which the DNA content is higher (x-y: z aspect ratio decreases). Intrinsic fluorescence gets progressively stronger with decreasing N:P, indicating detection of higher DNA content in the droplets formed at lower N:P. (**b**) Confocal intrinsic fluorescence images of PLL_240_-PRE_20_ droplets at increasing N:P in the x-y plane (top panels) z-y plane (bottom panels). Droplets are more elevated in the z-axis at lower N:P at which the PRE_20_ content is higher (x-y: z aspect ratio decreases). Intrinsic fluorescence is significantly weaker for PLL_240_-PRE_20_ despite comparable charge stoichiometry and concentration.

To test whether this is observed in non-DNA condensates, we studied the LLPS of PLL_240_ in presence of the negatively charged polypeptide, poly(R-glutamic acid) (PRE_20_), with chain length comparable to that of the DNA oligomer. **Figure 3b** shows droplets of PLL_240_-PRE_20_, formed at N:P ratios ranging from 0.2 to 3.5 (in this case, P= negatively charged amino acid residues of PRE). Circular droplets of PLL_240_-PRE_20_ are formed at N:P= 0.1 to 1.8 with the fluorescence intensity of the droplets remaining constant upon increasing N:P, unlike for PLL_240_-poly(A)_22_ droplets. However, the droplet contact angle for PLL_240_-PRE_20_ also decrease with increasing N:P (**Figure 3b bottom panel**, x-z plane), indicating the overall charge ratio of components undergoing LLPS is critical in dictating the resulting condensate morphology.

### Core-shell and multiphase architecture tuned is by charge ratio and DNA hybridization

In vitro studies revealing how LLPS of different proteins and nucleic acids can be tuned to yield different multiphase architectures can give us critical insights into the determinants of sub-compartmentalization in biomolecular condensates/MOs in cells. Using fluorescently tagged biomolecules, we and others have shown that hybridization can tune the phase behavior of DNA-based condensates (18, 24). With polypeptides like PLL, ssDNA favors droplet formation, while double stranded DNA favors precipitation. Here, we show that DNA hybridization in conjunction with varying N:P ratio leads to formation of hollow, membrane-like organization of DNA droplets displaying varying morphology. As we examine the intrinsic condensate fluorescence rather than that of fluorescent tags, the obtained images show direct evidence of spatiotemporal organization of the condensates. By utilizing the differences in intrinsic fluorescence intensities between DNA and non-DNA condensates, we can also reveal coexistence of multiple condensate phases.

To demonstrate the role of DNA hybridization, we added complementary ssDNA, poly(T)_22,_ to a series of preformed droplets of PLL_240_-poly(A)_22_ obtained at N:P ratios below 1 (containing higher DNA content) (**Figure 4**). Intrinsic fluorescence confocal video-microscopy shows immediate appearance of growing vacuoles inside the droplets (**Movies S1-4**). The vacuoles are slower to emerge in droplets formed at higher N:P ratio. The resulting condensates have a core-shell architecture with hollow interior as evidenced from 2D and 3D confocal images which show intrinsic fluorescence only at the condensate surface, while the interior does not fluoresce (**Figure 4b, S2)**. This demonstrates that all macromolecules concentrate at the surface of the core-shell condensates. The ability to form core-shell structure decreases as the N:P increase and at N:P > 1, the droplets have a coarser appearance, not a hollow interior. The z-profiles show the droplets become progressively flatter as the N:P ratio increases (**Figure 4b, bottom panel**). In absence of DNA hybridization, the core-shell morphology is not observed as indicated by homogenous intrinsic fluorescence from PLL_240_-poly(A)_22_ droplets after adding equimolar amounts of poly(A)_22_ (**Figure S3**).

**Figure 4.**
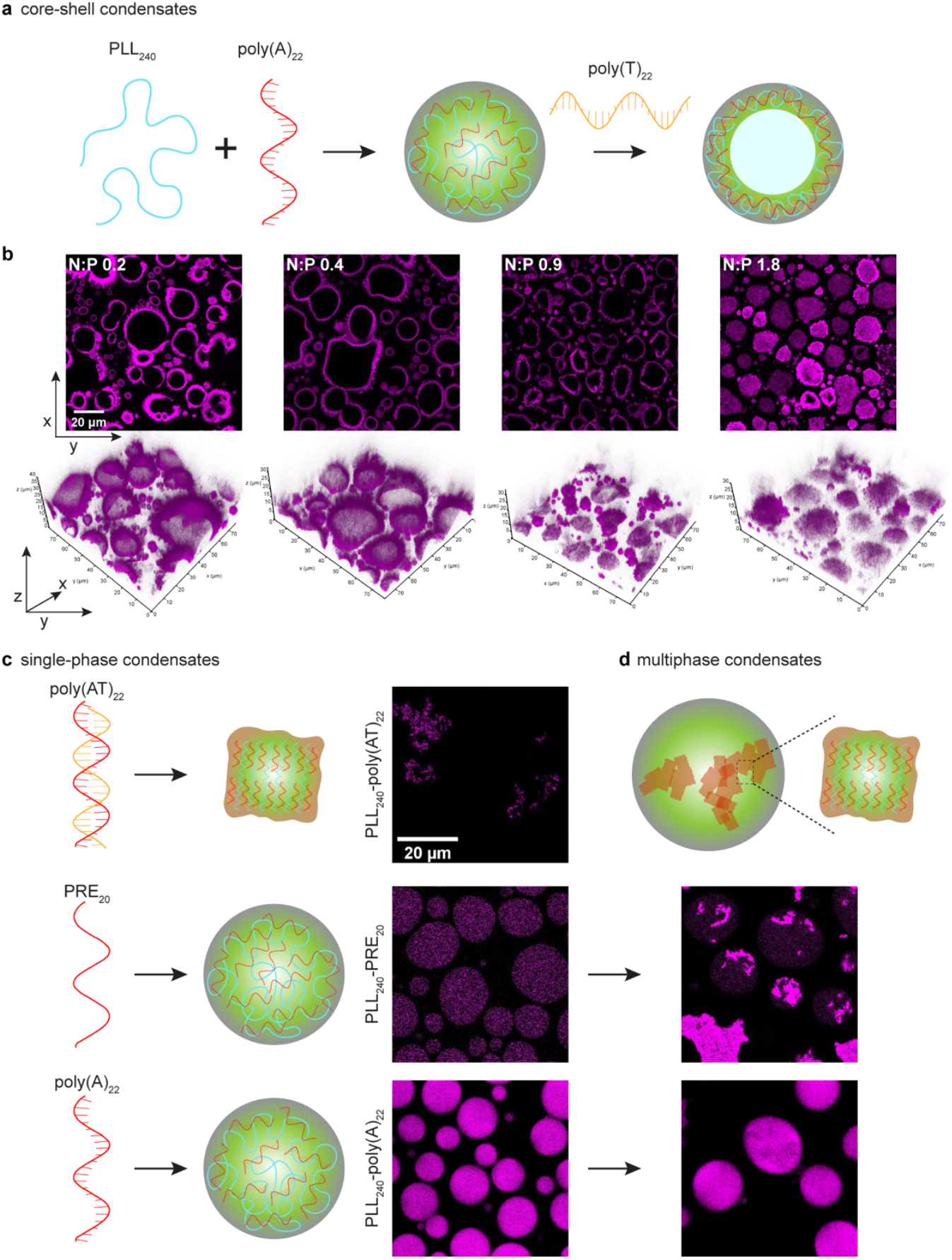
Core-shell and multiphase architecture of condensates is tuned by charge ratio and DNA hybridization. (**a**) Schematic of DNA hybridization driven core-shell multiphase architecture in pre-formed polypeptide-single stranded (ss) DNA droplets (**b**) Confocal intrinsic fluorescence images of PLL_240_-poly(A)_22_ droplets at increasing N:P after adding equimolar amounts of the complementary strand, poly(T)_22_ in 2D (top panels) and 3D (bottom panels). At low N:P, the droplets become hollow within seconds after adding the complementary strand (see **Figure S2**). At high N:P (> 1.0), there is higher propensity of gelation compared to formation of hollow condensates. 3D confocal images show that the condensate height is lower at higher N:P, consistent with that of droplets prior to hybridization-driven core-shell assembly. (**c**) Schematic of formation of precipitates of double stranded (ds) DNA and partitioning of the precipitates in pre-formed droplets to form multiphase condensate architecture. (**d**) Partitioning of precipitates of PLL_240_-poly(AT)_22_ (top panel) into pre-formed PLL_240_-PRE_20_ and PLL_240_-poly(A)_22_ droplets (N:P ∼1, buffer: 10 mM Tris-HCl buffer at pH 7.4 with 1 mM EDTA, 150 mM NaCl) resulting into multiphase droplets.

Previously, we have shown that DNA rigidity can tune LLPS of protein/polypeptide-DNA droplets. While dsDNA forms solid aggregates with charged polypeptides, ssDNA robustly forms liquid droplets (15, 18). Here, we study whether partitioning of solid condensates into preformed liquid-like condensate occurs, and, if they result in multiphase condensates (**Figure c-d**). As described above (**Figure 1-2**), intrinsic fluorescence is detected in both droplets with and without DNA. Notably, intrinsic fluorescence from droplets containing DNA is significantly higher. This relative difference in intrinsic fluorescence can be utilized to image multi-phase condensates. **Figure 4d** shows that precipitates of PLL_240_-poly(AT)_22_ (top panel), which, when mixed with either PLL_240_ -PRE_20_ (middle panel) or PLL_240_-poly(A)_22_ droplets (bottom panel), are engulfed by both types of droplets and coexist as a separate phase. As the intrinsic fluorescence of PLL_240_ -PRE_20_ droplets are weaker in intensity compared to PLL_240_-poly(AT)_22_ precipitates, they appear as a bright inhomogeneous phase inside PLL_240_ -PRE_20_ droplets. Conversely, as the intrinsic fluorescence from the PLL_240_-poly(A)_22_ droplets are higher in intensity compared to PLL_240_-poly(AT)_22_ precipitates, they appear as dark inhomogeneous phase inside PLL_240_-poly(A)_22_ droplets. Such relative intensity differences in intrinsic fluorescence from different condensates can be utilized to directly assess different phases in multiphase condensates to determine various factors that affect the formation and maintenance.

## Discussion

Due to the potential connection of cellular organization and myriad human diseases, membraneless organelle and biomolecular condensate formation and regulation in cells has become a topic of substantial importance (2-7). There is still a long way to go in understanding the formation of biomolecular condensates. These condensates often display intricate multilayered architectures inside the cell for example, the nucleolus (25) and nuclear speckles (26). An increasing number of proteins are known to concentrate in biomolecular condensates via phase separation. The most common identifying features of proteins that tend to undergo LLPS include high content of charged residues, IDRs (intrinsically disordered regions), and modular domains (27, 28). This has led to a number of growing databases dedicated to predicting the phase separation propensity of proteins (29, 30). Consideration should be taken as there is experimental evidence indicating that protein phase behavior cannot always be predicted simply by applying common structural feature-based methods (16, 31). Most often proteins have different RNA/DNA binding partners that facilitate the formation of biomolecular condensates(2, 32). This vast scope of protein phase separation behavior necessitates reliable and efficient methods to examine large sets of data on the phase separation of proteins and improve the prediction of protein/NA phase transitions and multiphase architectures.

Owing to their high sensitivity and selectivity, fluorescence-based methods that employ engineered fluorophore tags are currently the predominant method for detecting biomolecular condensates(8, 9). There is always a concern fluorophore-tagged proteins are not representative of the native protein structure, or that interactions with the tags have unknown contributions to condensation (8). For in vitro labeled samples, incomplete purification of tagged proteins and NAs can result in free fluorescent chromophores causing background noise that interferes with data interpretation. Labeling techniques are also often time consuming and expensive. These aspects limit high throughput analysis of protein/NA phase separation based on external fluorophore labeling thus necessitating development of label-free techniques for efficient experimental analysis of phase behaviors of proteins.

To this end, we leveraged the intrinsic fluorescence arising as an emergent behavior of condensates, possibly via AIE or related mechanisms (7). We detected LLPS of histones in presence of a variety of G-quadruplexes. Not only is the technique presented here useful for efficient detection of condensate formation, but also allows us to measure condensate dynamics via monitored merging events and FRAP. We were able to demonstrate modulation of condensate morphology and multi-phase architecture and showed that by varying concentration ratio of positive and negative species, the 3D morphology of condensates can be tuned through modulation in overall charge balance (as represented by N:P ratio). Although this concept is commonplace in cationic polymer mediated NA delivery for nucleic acid therapy-based studies (33-35), the charge ratio concept is often ignored in studies of biomolecular condensates. Our study directly demonstrates the degree of overall charge balance between scaffold molecules is a complementary approach to predicting both phase behavior and morphology of protein-NA condensates. We also demonstrate NA hybridization driven multiphase behavior, also tunable via the N:P ratio. Formation of core-shell condensates have been previously demonstrated by us and others in NA-containing condensates using fluorophore tagged components (18, 36). Here, we directly demonstrate that under the conditions that drive formation of core-shell condensates, the scaffold components migrate to the surface resulting in membrane-like exterior.

Our method takes advantage of emergent optical properties resulting from the high concentrations of proteins/NAs/polymers within condensates and can easily be applied to high-throughput studies. This approach can greatly advance the development of a general framework to describe the features of proteins/NA controlling condensate architectures. Localization of individual molecules in the absence of molecule-specific fluorescence tags cannot be easily assessed using intrinsic fluorescence, we however find that intensity differences can help overcome this limitation to a degree for multicomponent systems of both proteins and nucleic acids. By utilizing the differences in emission intensities between protein-rich condensates vs NA-rich condensates, partitioning of solid-like condensates inside liquid-like condensates can be studied. Further studies characterizing the optical properties of condensates such as excitation wavelength dependence of emission and resolving lifetimes can, to some extent, help overcome the limitation of specificity of intrinsic fluorescence emission-based imaging. In principle, intrinsic fluorescence-based condensate imaging can be incorporated in super-resolution fluorescence and fast confocal imaging modules.

In summary, our findings, through use of a novel label-free fluorescence method that detects a fundamental property of biomolecular condensates, reveal various factors that drive biomolecular condensate formation and their multiphase organizations. This method may facilitate the development of new high-throughput techniques in this quickly evolving and expanding field of study into intracellular LLPS. Our findings open a new avenue towards understanding the complex phase transitions driven membraneless organization of biomolecules in cells.

## Materials and Methods

### Materials

All DNA oligomers (quadruplex and linear DNA sequences) were obtained from Integrated DNA Technologies (Skokie, IL). The sequences of oligomers are provided in supplementary information (**SI Table 1**). Histone H1 was obtained from Sigma-Aldrich (St. Louis, MO). PRE_20_, PLE_100_, PLD_100_, and PRK_100_ were obtained from Alamanda Polymers (Huntsville, AL). PLL_240_ (MW 30-70 kDa), Tris-EDTA buffer (10 mM Tris-HCl, 1 mM EDTA, pH 7.4), and sodium chloride were obtained from Sigma (St. Louis, MO).

### DNA, polypeptide, and protein sample preparation

All oligonucleotide stock solutions, except G-quadruplexes, were prepared by dissolving in the above Tris-EDTA buffer (10 mM Tris-HCl, 1 mM EDTA, pH 7.4) without added salt and annealing (at ∼2 mM oligomer concentration) by heating at 95°C for 5 minutes and cooling for at least 1 hr. The concentrations were measured using NanoDrop (ThermoFischerScientific) and samples were tested for size and purity using gel electrophoresis. G-quadruplexes were prepared in 10 mM potassium phosphate buffer with 100 mM KCl, at pH=7.4. Folding topology of the G-quadruplexes were verified using circular dichroism CD spectroscopy (**Fig. 1d**). All polypeptide stock solutions were prepared by dissolving in H_2_O at target concentration of 20 mM. Histone H1 stock solution was prepared in Tris-EDTA buffer (10 mM Tris-HCl, 1 mM EDTA, pH 7.4) at 1 mM concentration.

### Condensate preparation

Eight-well borosilicate (1.0) chambered glass slides (Nunc Lab Tek; ThermoFischerScientific) were cleaned with RNase Zap (Ambion; ThermoFischerScientific), ethanol, and nuclease-free water. The glass surface was passivated using 3.5% bovine serum albumin solution (Sigma) for 15 min at room temperature, rinsed three times with nuclease-free water, and dried. All sample preparation and imaging was performed at room temperature.

For the preparation of condensates (droplets/precipitates), the stock solutions of all negatively charged macromolecules (DNA and the polypeptides PRE_20_, PLE_100_, and PLD_100_) were diluted to desired concentrations in the Tris-EDTA buffer without added salt. Stock solutions of all corresponding positively charged macromolecules (polypeptides PRK_100_, and PLL_240_) were diluted to desired concentrations in the Tris-EDTA buffer containing added NaCl. For H1-G-quadruplex droplets, the same method was followed except G-quadruplex folding buffer was used (10 mM phosphate buffer with 50 mM KCl). The dilutions of positively charged species and negatively charged species were mixed in 1:1 volume, such that the final N:P (ratio of net positive charge to net negative charge in solution) was ∼1. The final salt concentration was 150 mM NaCl except for H1-G-quadruplex droplets (50 mM KCl). The order of mixing of solutions was always positively charged protein/polypeptide added to negative charged DNA/polypeptide solution. The resulting mixture was then added to the passivated eight-well chambers for subsequent bright-field, confocal intrinsic fluorescence, or FRAP imaging as described below.

### Bright-field imaging

Bright-field images were obtained using a Leica DMI6000 B microscope (Leica Microsystems, Wetzlar, Germany) using 100X oil objective and Grasshopper3 camera (Point Grey, Richmond, BC, Canada). Images were processed to adjust brightness, contrast, and image size using the ImageJ software (37).

Bright-field images of drops and precipitates were taken ∼2 hours after mixing the positively and negatively charged polymer solutions. For droplet-precipitate mixing experiments, the 2-hour precipitates were added to the 2-hour pre-formed droplets in the passivated chambers. The mixing of drop precipitate was monitored by taking images at a 2-hour and 24-hour time point.

### Confocal imaging

All 2D and 3D confocal microscopy and FRAP was carried out on a Leica SP8X microscope (Leica microsystems) equipped with pulsed laser excitation (NKT Photonics, Birkerod, Denmark), single-photon avalanche diodes (SPADs) (Micron Photon Devices, Bolzano, Italy), and external time-correlated single photon counting (TCSPC) detection (PicoQuant, Berlin, Germany).

### Native fluorescence imaging

For confocal imaging using native fluorescence, excitation was carried out at 532 nm (white light laser at 50% intensity and 70% master power). Emission was filtered from 550-750 nm and detected on a SPAD detector. The excitation spectrum was collected using a set emission detection bandwidth (550-750 nm) as the excitation was systematically scanned between 472-532 nm. Detection was carried out on a SPAD detector. The emission spectrum was collected using a set 532 nm excitation and detecting emission from 10 nm bandwidth windows with a center wavelength systematically varied from 560 nm to 690 nm. Detection was carried out on a HyD detector.

### Fluorescence recovery after photobleaching (FRAP)

FRAP was carried out using 532 nm excitation (white light laser at 50% intensity and 70% master power) for imaging and 532 nm excitation (white light laser at 100% intensity and 70% master power) for bleaching. Bleaching was done at a single region of interest for 1 s. Imaging was carried out at ∼3 fps, and recovery was measured for over 100 s.

### CD spectroscopy

Circular dichroism (CD) spectra of G-quadruplexes were obtained using JASCO J-815 spectropolarimeter (JASCO, Inc., Japan). The instrument was subjected to a 15-min nitrogen purge before activating the light source, establishing an oxygen-free environment to prevent UV absorption by atmospheric oxygen. Throughout the experimental duration, a continuous nitrogen flow was maintained to uphold the inert atmosphere. After the nitrogen purge, the lamp power was initiated, and the Peltier device was activated. To ensure temperature stability, a water circulator facilitated controlled water circulation, maintaining the experimental temperature at 20°C. A stabilization period of approximately 30 min was observed to allow the light source to reach equilibrium.

A quartz cuvette with a width of 10 mm was used to load the sample. 5μM of G-quadruplex samples in annealed in folding buffer were used. For the control unfolded samples, NaOH was added to the buffer. CD spectra readings were acquired in the range of 340-200 nm at a scanning speed of 50 nm/min, a data integration time of 1 s, a data pitch of 0.5 nm, and bandwidth of 1 nm. All the parameters were set using the SpectraManager 2 graphical user interface. For each sample, five spectra data sets were collected, and the cuvette was rinsed with DI water, followed by ethanol, and dried using N_2_ gas after each sample change. Baseline measurements were recorded, and manual baseline correction was performed.

