## Supporting Info for "Label-Free Fluorescence Microscopy Reveals Multiphase Organization in Biomolecular Condensates"

**This PDF file includes:**

Figures S1 to S3
Tables S1
Legends for Movies S1 to S4
SI References

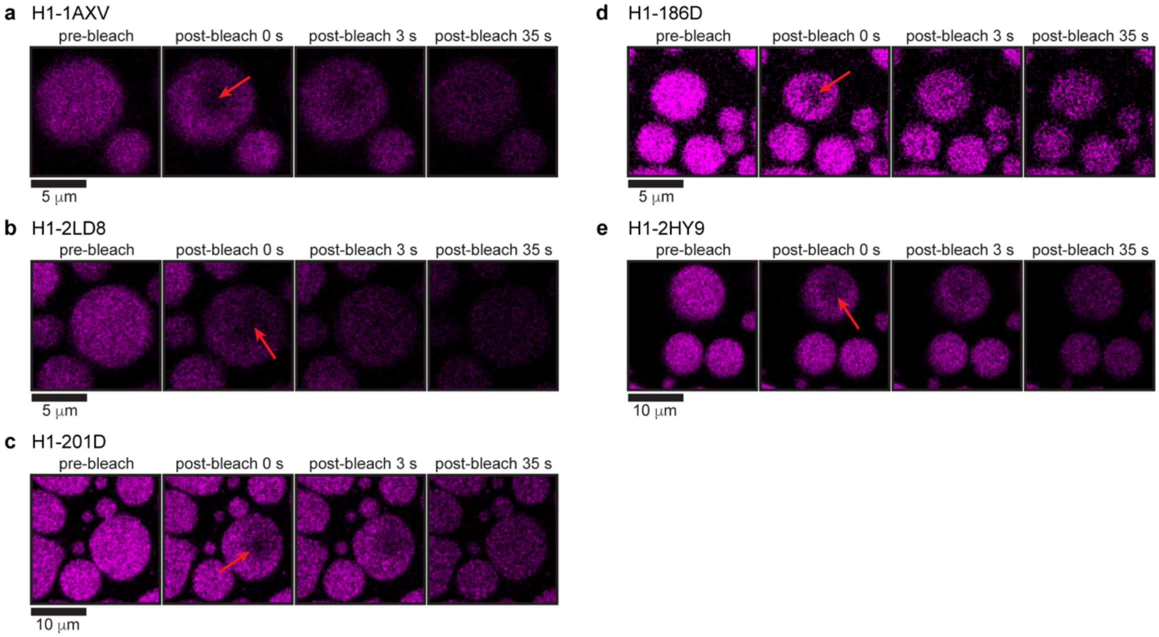


**Figure S1.** Intrinsic FRAP in condensates of histone H1 with five different G-quadruplex DNA (**a**-**e**), showing pre-bleach images and fluorescence recovery images at three different time points (0, 3, and 35 s).

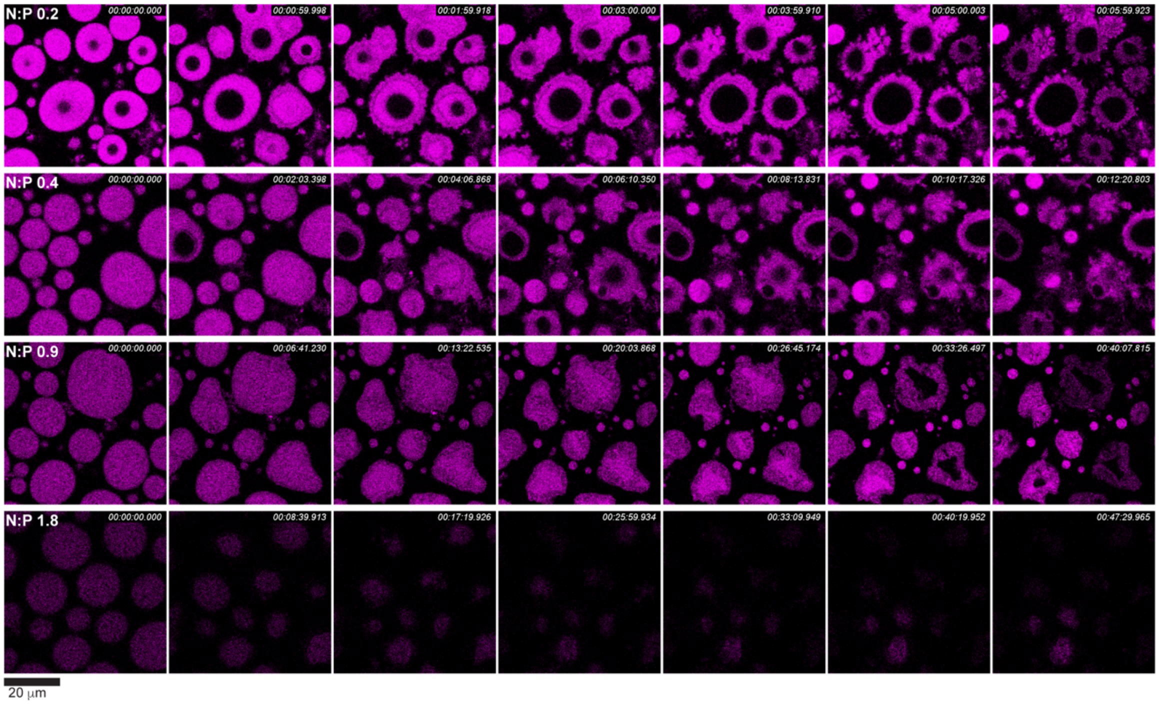


**Figure S2.** Time series confocal intrinsic fluorescence images of $PLL_{240}$-poly(A)$_{22}$ droplets at increasing N:P after adding equimolar amounts of poly(T)$_{22}$. Formation of vacuoles inside the droplets is significantly slows down as N:P increases with no core-shell formation at N:P > 1.

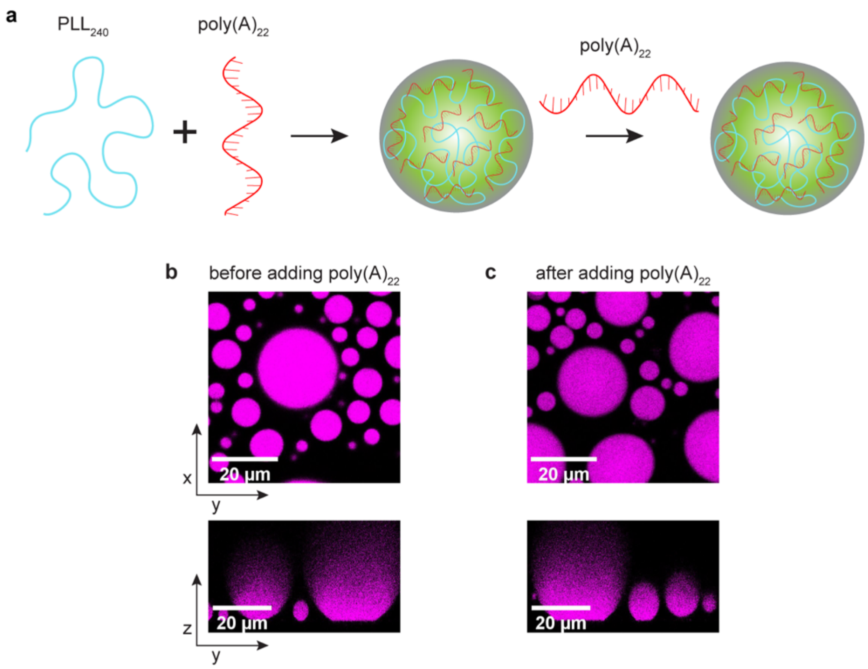


**Figure S3.** (**a**) Schematic of the morphology of pre-formed $PLL_{240}$-poly$(A)_{22}$ droplets after adding poly$(A)_{22}$. Confocal intrinsic fluorescence images of $PLL_{240}$-poly$(A)_{22}$ droplets in x-y and z-y planes before (**b**) and after (**c**) adding equimolar amounts of poly$(A)_{22}$ to pre-formed droplets formed at N:P 0.2.

**Table S1.** Sequences of ssDNA oligomers and G-quadruplex DNA used in this study.

| | **Sequence** | **Length (nt)** |
|---|---|---|
| ***G-quadruplexes***[1] | | |
| 1AXV | 5'-TGAGGGTGGGTAGGGTGGGTAA | 22 |
| 2LD8 | 5'-TAGGGTTAGGGTTAGGGTTAGGG | 23 |
| 201D | 5'-GGGGTTTTGGGGTTTTGGGGTTTTGGGG | 28 |
| 186D | 5'-TTGGGGTTGGGGTTGGGGTTGGGG | 24 |
| 2HY9 | 5'-AAAGGGTTAGGGTTAGGGTTAGGGAA | 26 |
| ***DNA oligos*** | | |
| poly(A)$_{22}$ | 5'-AAAAAAAAAAAAAAAAAAAAAA | 22 |
| poly(T)$_{22}$ | 5'-TTTTTTTTTTTTTTTTTTTTTT | 22 |
| poly(AT)$_{22}$ | 5'-AAAAAAAAAAATTTTTTTTTTT | 22 |

**Movie S1 (separate file).** Intrinsic fluorescence confocal video-microscopy of $PLL_{240}$-poly$(A)_{22}$ droplets formed at N:P 0.2 after adding equimolar amounts of poly$(T)_{22}$.

**Movie S2 (separate file).** Intrinsic fluorescence confocal video-microscopy of $PLL_{240}$-poly$(A)_{22}$ droplets formed at N:P 0.4 after adding equimolar amounts of poly$(T)_{22}$.

**Movie S3 (separate file).** Intrinsic fluorescence confocal video-microscopy of $PLL_{240}$-poly$(A)_{22}$ droplets formed at N:P 0.9 after adding equimolar amounts of poly$(T)_{22}$.

**Movie S4 (separate file).** Intrinsic fluorescence confocal video-microscopy of $PLL_{240}$-poly$(A)_{22}$ droplets formed at N:P 1.8 after adding equimolar amounts of poly$(T)_{22}$.
